# Proteins in the periplasmic space and outer membrane vesicles of *Rhizobium etli* CE3 grown in minimal medium are largely distinct and change with growth phase

**DOI:** 10.1101/305797

**Authors:** Hermenegildo Taboada, Niurka Meneses, Michael F. Dunn, Carmen Vargas-Lagunas, Natasha Buchs, Jaime A. Castro-Mondragon, Manfred Heller, Sergio Encarnación

## Abstract

*Rhizobium etli* CE3 grown in succinate-ammonium minimal medium (MM) excreted outer membrane vesicles (OMVs) with diameters of 40 to 100 nm. Proteins from the OMVs and the periplasmic space were isolated from 6 and 24 h cultures and identified by proteome analysis. A total 770 proteins were identified: 73.8 and 21.3 % of these proteins occurred only in the periplasm and OMVs, respectively, and only 4.9 % were found in both locations. The majority of proteins found in either location were present only at 6 or 24 h: in the periplasm and OMVs, only 24 and 9 % of proteins, respectively, were present at both sampling times, indicating a time-dependent differential sorting of proteins into the two compartments. The OMVs contained proteins with physiologically varied roles, including *Rhizobium* adhering proteins (Rap), polysaccharidases, polysaccharide export proteins, autoaggregation and adherence proteins, glycosyl transferases, peptidoglycan binding and cross-linking enzymes, potential cell wall modifying enzymes, porins, multidrug efflux RND family proteins, ABC transporter proteins, and heat shock proteins. As expected, proteins with known periplasmic localizations (phosphatases, phosphodiesterases, pyrophosphatases) were found only in the periplasm, along with numerous proteins involved in amino acid and carbohydrate metabolism and transport. Nearly one-quarter of the proteins present in the OMVs were also found in our previous analysis of the *R. etli* total exproteome of MM-grown cells, indicating that these nanoparticles are an important mechanism for protein excretion in this species.

**IMPORTANCE:** The reduction of atmospheric nitrogen to ammonia by rhizobia symbiotically associated with legumes is of major importance in sustainable agricultural. Rhizobia excrete a variety of symbiotically important proteins using canonical secretion systems. In this work, we show that *Rhizobium etli* grown in culture also excretes proteins in membrane-enclosed structures called outer membrane vesicles (OMVs). This study reports OMV production by rhizobia. Proteins identified in the OMVs included Rhizobium adhering (Rap) and autoaggregation proteins, polysaccharidases, RTX toxins, porins and multidrug efflux proteins. Some of these proteins have important roles in the *R. etli*-common bean symbiosis, and their packaging into OMVs could deliver them to the environment in a concentrated yet diffusible form protected from degradation. The work described here provides a basis for future studies on the function of rhizobial OMVs in free life and symbiosis.

## INTRODUCTION

Bacterial protein secretion is a vital function involving the transport of proteins from the cytoplasm to other cellular locations, the environment, or to eukaryotic host cells (1). Of the proteins synthesized by *Escherichia coli* on cytoplasmic ribosomes, about 22 % are inserted into the inner membrane (IM), while 15 % is targeted to periplasmic, outer membrane (OM) and extracellular locations (2).

The IM is a phospholipid bilayer that surrounds the cytoplasm. The OM is comprised of an inner leaflet containing phospholipids and lipoproteins and an outer leaflet comprised mostly of lipopolysaccharide (LPS) and also containing proteins such as porins (3). The periplasmic space of gram-negative bacteria is delineated by the IM and OM, with a thin peptidoglycan layer attached to both membranes by membrane anchored proteins. The periplasm of *E. coli*, for example, contains hundreds of proteins including transporters, chaperones, detoxification proteins, proteases and nucleases (4, 5).

About a dozen specialized export systems for bacterial protein secretion have been described (1). Gram-negative bacteria also excrete proteins and other substances in outer membrane vesicles (OMVs). Phospholipid accumulation in the OM triggers the formation of these spherical structures, which are composed of a membrane bilayer derived from the bacterial OM (6). The amount of OMVs produced by a given bacterium varies in response to environmental conditions like growth phase, nutrient sources, iron and oxygen availability, abiotic stress, presence of host cells, and during biofilm formation (7). Depending on the species and growth conditions, OMVs may enclose cytoplasmic, periplasmic and transport proteins, as well as DNA, RNA, and outer membrane-derived components such as LPS and phospholipids. The inclusion of proteins in the OMVs is not random but appears to be determined by specific sorting mechanisms (2–5, 8). Suggested roles for OMVs include invasion, adherence, virulence, antibiotic resistance, modulation of the host immune response, biofilm formation, intra and interspecies molecule delivery, nutrient acquisition, and signaling (2, 8–12).

Rhizobia are gram-negative bacteria that reduce atmospheric nitrogen to ammonia in symbiotic association with leguminous plants. The excretion of specific proteins and polysaccharides by rhizobia is an essential component of this process (13). The alpha-proteobacterium *Rhizobium etli* CE3 establishes a nitrogen-fixing symbiosis with *Phaseolus vulgaris* (common bean). We have shown that *R. etli* CE3 secretes many proteins during exponential and stationary phase growth in minimal medium cultures (14) and suggested that some of the secreted proteins might be exported in OMVs, although these nanoparticles have not been reported in rhizobia (15). Mashburn-Warren and Whiteley have hypothesized that hydrophobic rhizobial nodulation (Nod) factors could be packaged in OMVs for delivery to the plant root, where they induce plant responses required for nodulation (10). OMVs produced by symbiotic rhizobia have not, however, been studied experimentally.

Relatively few studies have been done on OMVs produced by plant-associated bacteria, where these protein and molecule-bearing structures could enhance the benefits obtained by the prokaryote in mutualistic or pathogenic interactions (16, 17). Our major aim in this work was to identify proteins present in purified *R. etli* OMVs obtained from cells grown in culture. Because periplasmic proteins could be (perhaps nonspecifically) incorporated into the OMVs, we also identified proteins in the periplasm of cells grown under the same conditions. A major finding was that only a small fraction of the periplasmic proteins were also present in OMVs, which suggests that they are not randomly incorporated into the latter during vesicle formation. Our data also indicate that nearly one-quarter of the previously identified exoproteins produced by *R. etli* (15) are excreted in OMVs.

## RESULTS AND DISCUSSION

### General characteristics of the *R. etli* periplasmic and OMV fractions

We used proteome analysis to identify proteins in the periplasmic and OMV fractions prepared from *R. etli* cultures grown in MM for 6 and 24 h (Table S1, Supplemental material). Only proteins found in two experimental replicates are included in the dataset (see Materials and Methods). The suitability of hypo-osmotic shock protocols like the one used here to isolate periplasmic proteins has been demonstrated in several rhizobia (18, 19). The presence of cytoplasmic proteins in periplasmic protein preparations is commonly reported in the literature (see below) and could be an artifact resulting from cell lysis (20). In the *Pseudomonas aeruginosa* periplasm (obtained by spheroplasting), 39 and 19 % of 395 proteins identified were predicted to be cytoplasmic and periplasmic, respectively (21). The periplasmic proteomes of *Pseudoalteromonas haloplanktis* (22) and *Xanthomonas campestris* pv. *campestris* (23), both obtained by hypo-osmotic shock, contained many cytoplasmic enzymes for carbohydrate and amino acid metabolism, among others. In contrast, over 94 % of the 140 proteins obtained by osmotic shock and identified in the periplasms of *E. coli* strains BL21(DE3) and MG1655 were predicted to be periplasmic (24).

We used the PSLPred program to predict the subcellular localization of *R. etli* proteins. While the localization predictions made with this program are over 90 % accurate for proteins from gram negative bacteria (25), we noted that several proteins had an unexpected predicted localization. For example, the ribosomal proteins S12 and L31 were predicted to be periplasmic rather than cytoplasmic. Analysis of the *R. etli* exoproteome using an alternative protein localization prediction program, LocTree3 gave significantly different results in comparison to PSLpred. In the case of the ribosomal proteins described above, LocTree3 predicted that S12 is in fact cytoplasmic, but that L31 was secreted.

Based on PSLpred, the predicted cellular localizations of the 568 proteins found only in the *R. etli* periplasm (Table S2, Supplemental material) were 57 % cytoplasmic, 25 % periplasmic, 14.9 % IM, 1.8 % extracellular and 1.4 % OM. Importantly, the 8 highest-abundance proteins found in the total proteome of *R. etli* CE3 grown in MM (27) under conditions similar to those used in the present study were absent from the periplasmic or OMV fractions (Table S1, Supplemental material). This result argues against significant contamination of the periplasmic and OMV fractions with proteins resulting from cell lysis. Among the proteins identified in the *R. etli* OMVs and periplasm, all that are classed as phosphatases, phosphodiesterases or pyrophosphatases were found only in the periplasm (Table S1, Supplemental material), consistent with the biochemically determined localization of these enzymes in other rhizobia (18, 28). In addition, electron microscopic examination of the cell preparations obtained after hypo-osmotic shock showed that the IMs were still intact (results not shown).

The *R. etli* OMV fraction was obtained by differential centrifugation of culture filtrates. Transmission electron microscopic examination of the OMVs purified from 6 and 24 h cultures showed that the vesicles were spherical and had diameters of 40 to 100 nm, within size range expected for OMVs (16, 29). No pili, bacteria, flagella or membrane debris were detected (Fig. 1). SDS-PAGE analysis showed that the OMV protein patterns differed significantly from those of whole cell extracts (data not shown), consistent with the proposal that specific protein sorting mechanisms are important in determining the protein content of bacterial OMVs (3, 10, 30).

**Fig. 1.**
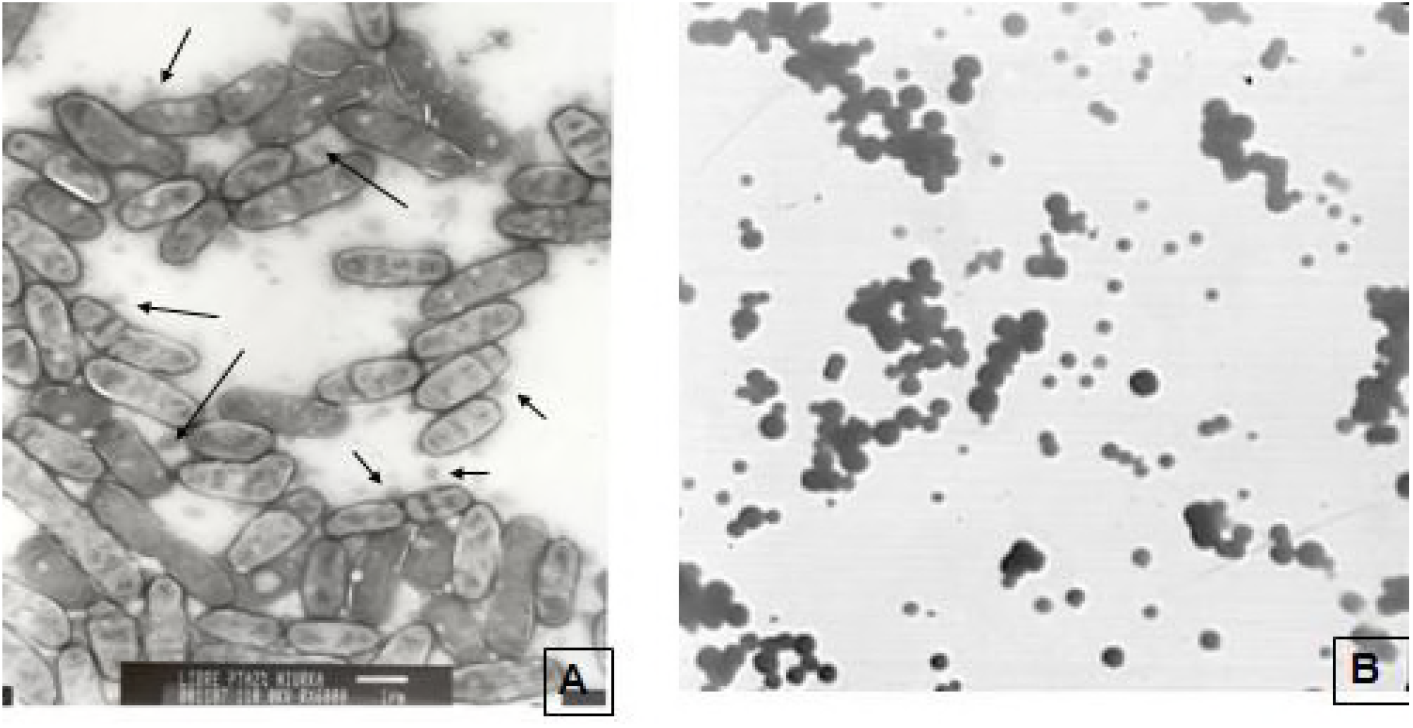
OMVs from *Rhizobium etli* CE3. Panel A. Electron micrograph of a culture of strain CE3 showing OMVs (arrows). Panel B. Electron micrograph of purified OMVs.

For proteins present only in the OMVs (Table 1), 39 and 34 % were predicted to be periplasmic and cytoplasmic, respectively, followed by 14 % IM, 8 % OM and 4 % extracellular. Although the presence of cytoplasmic and IM proteins as *bona fide* components of bacterial OMVs is controversial (3, 30), they have been found in similar proportions in OMVs from several species (29, 31–34). In comparison to the periplasm, the OMVs contain 5.7 times the number of OM proteins, consistent with the enrichment of these proteins in bacterial OMVs (30). For the 38 proteins found in both the periplasmic and OMV fractions (Table S3, Supplemental material), the predicted localizations were biased towards periplasmic proteins (53 %), followed by cytoplasmic (29 %), IM (16 %), OM (3 %) and extracellular proteins (0 %).

**Table 1.**
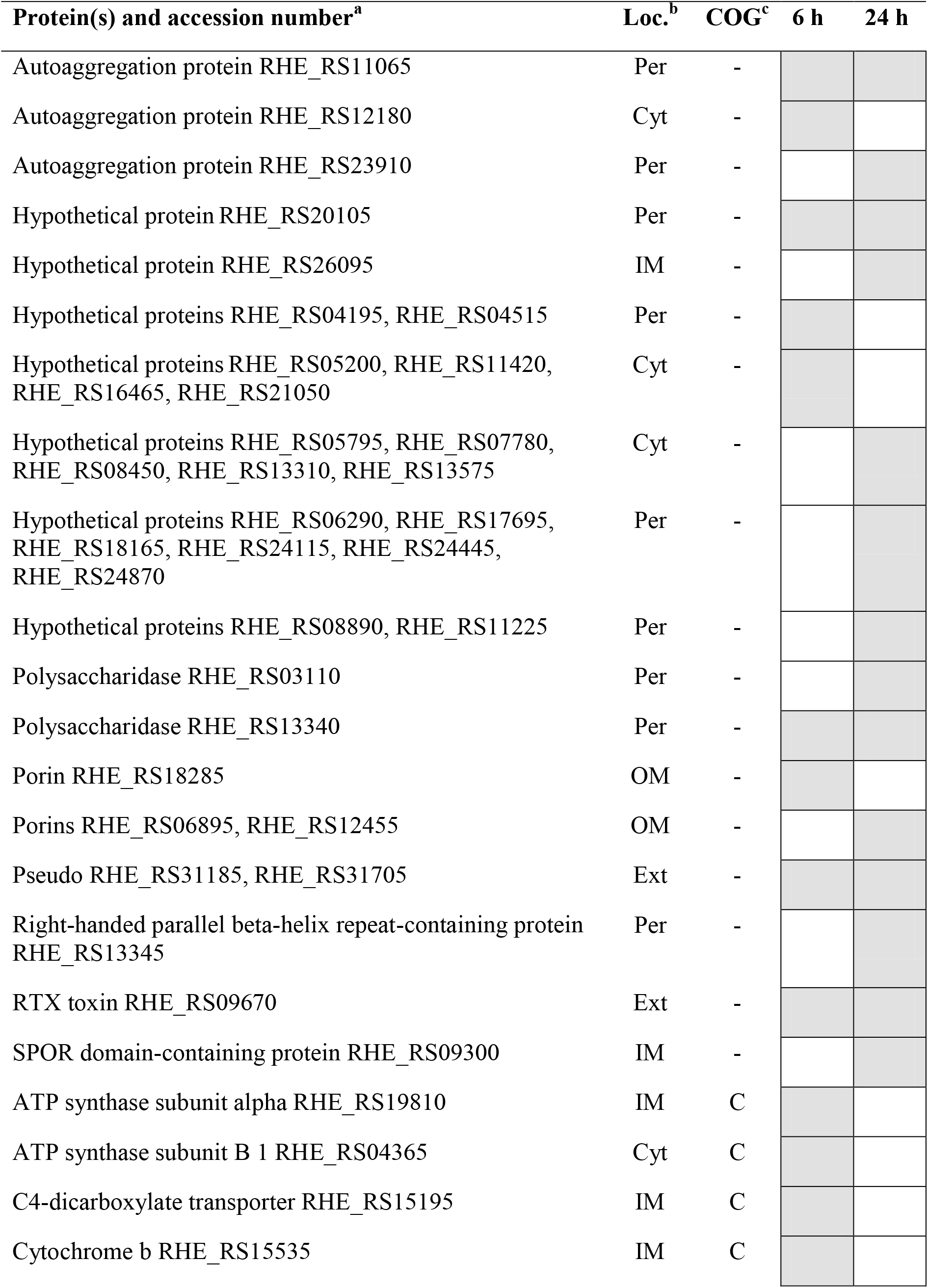
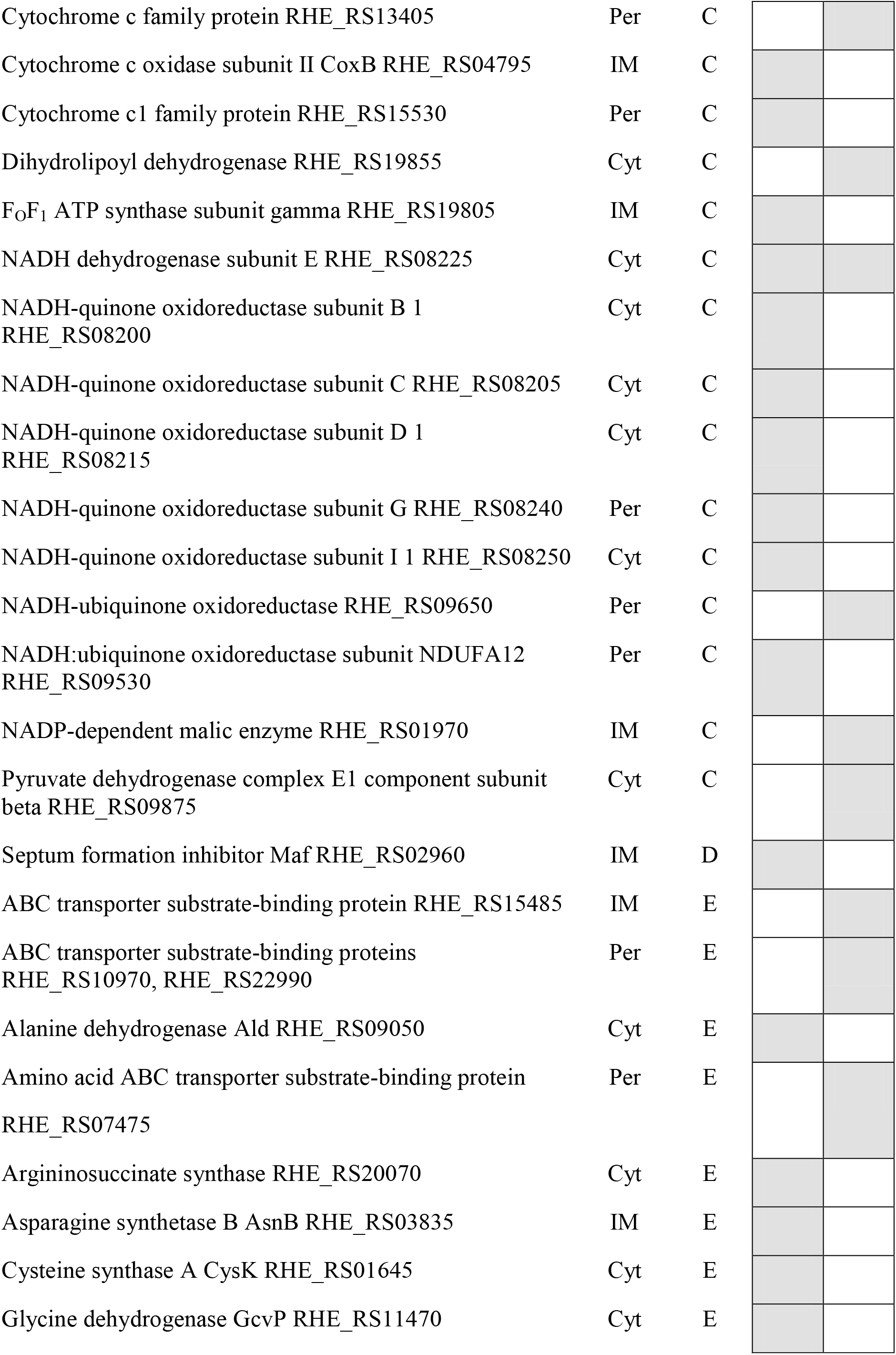
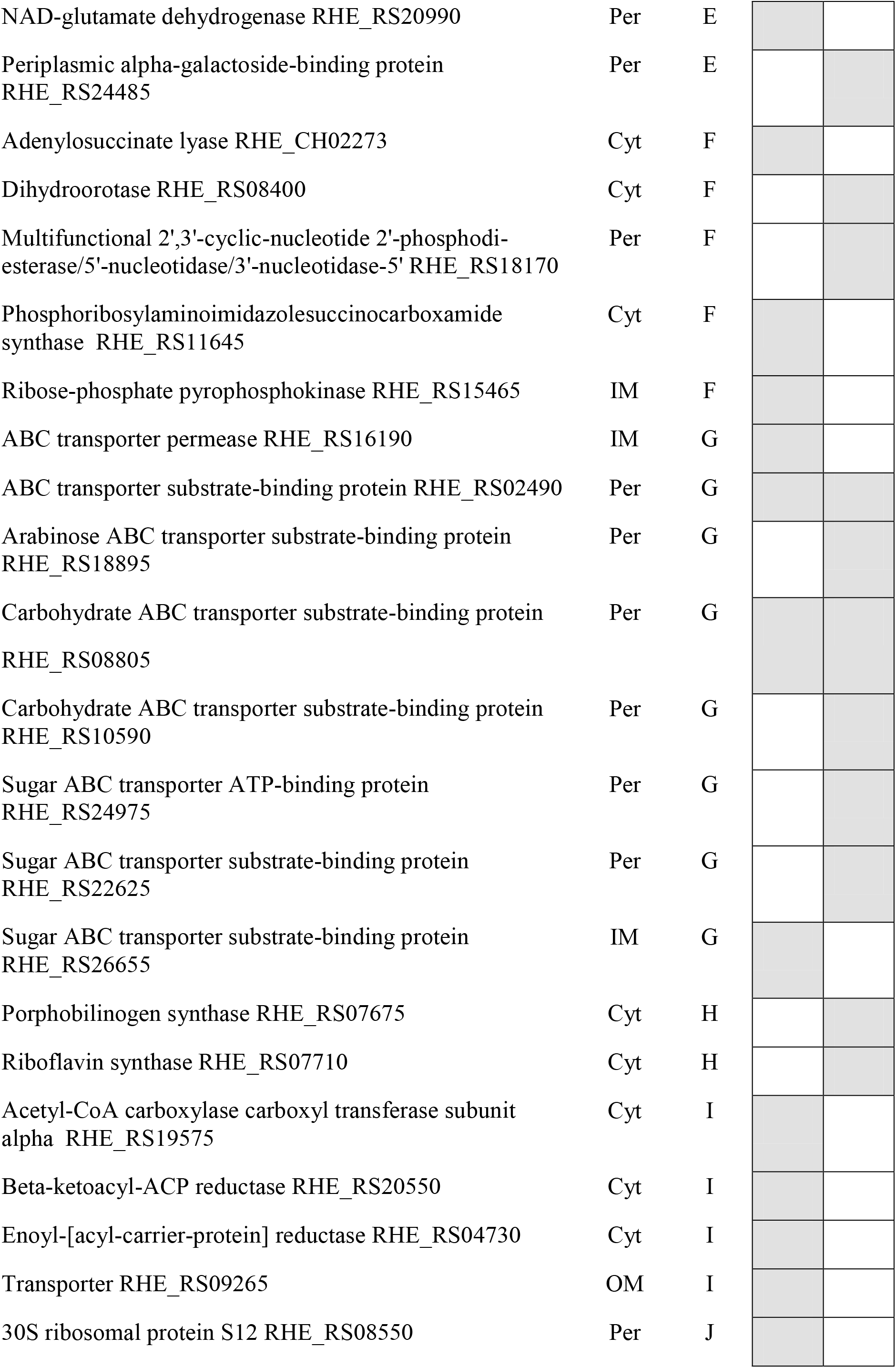
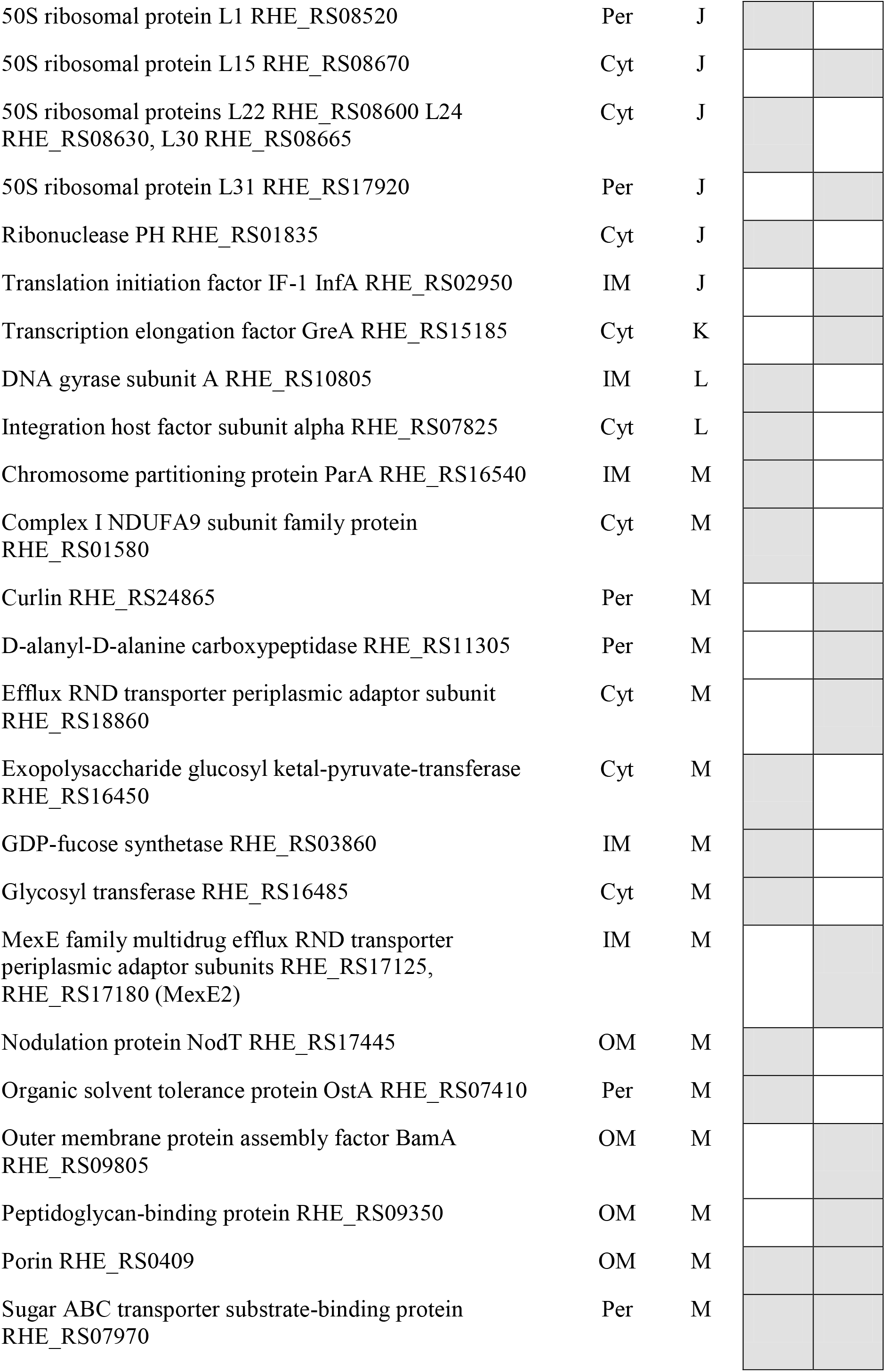
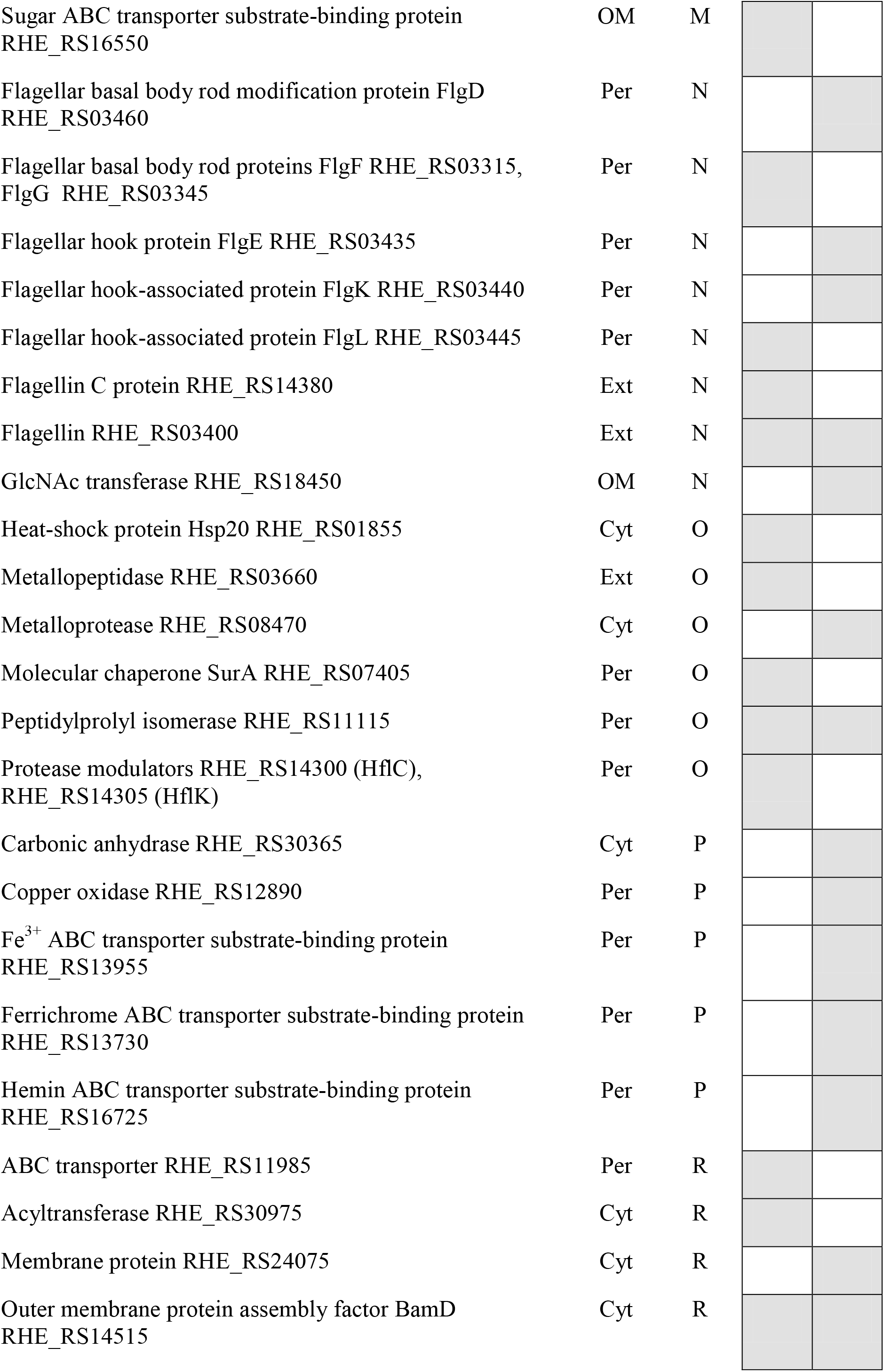
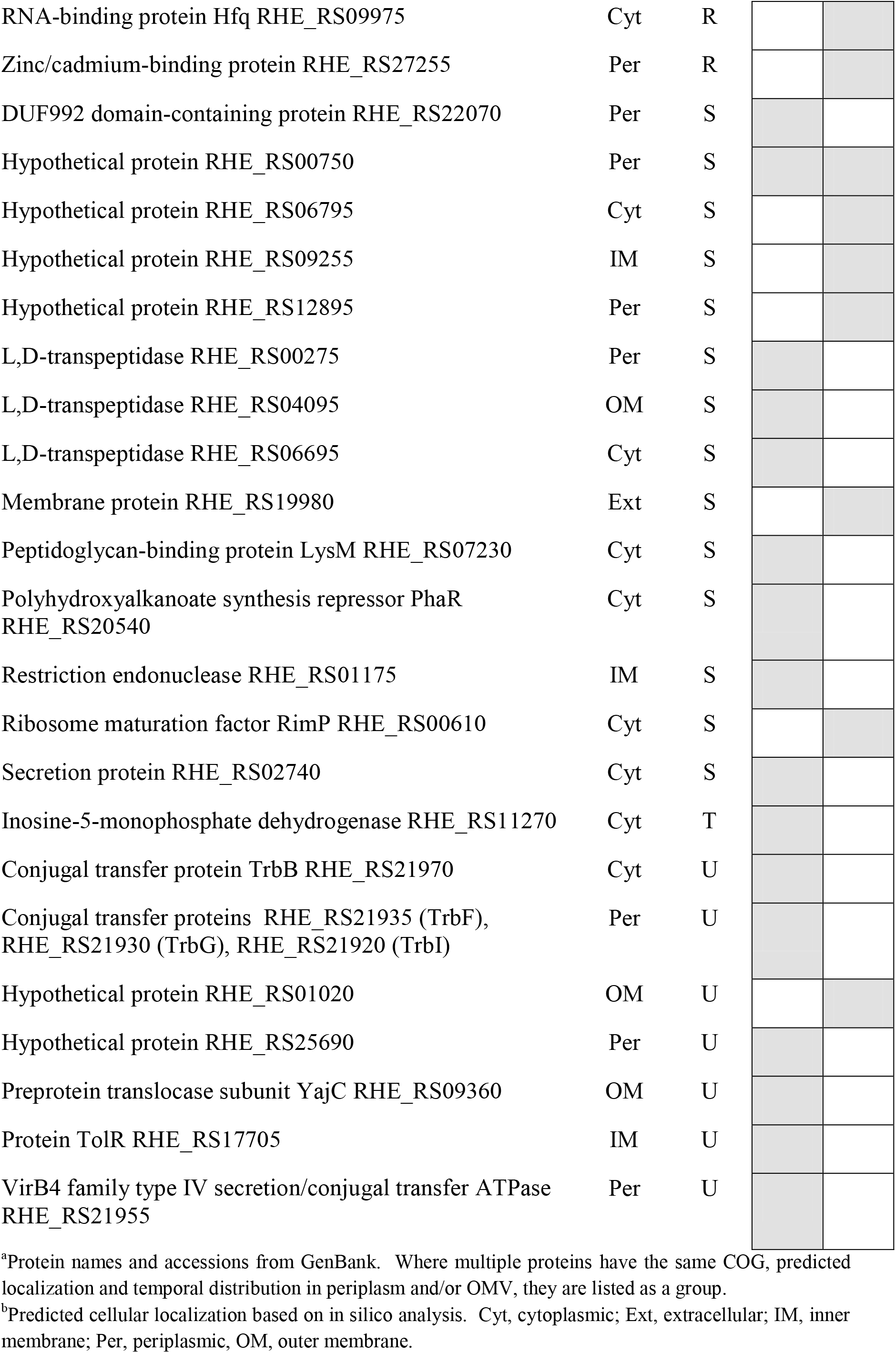
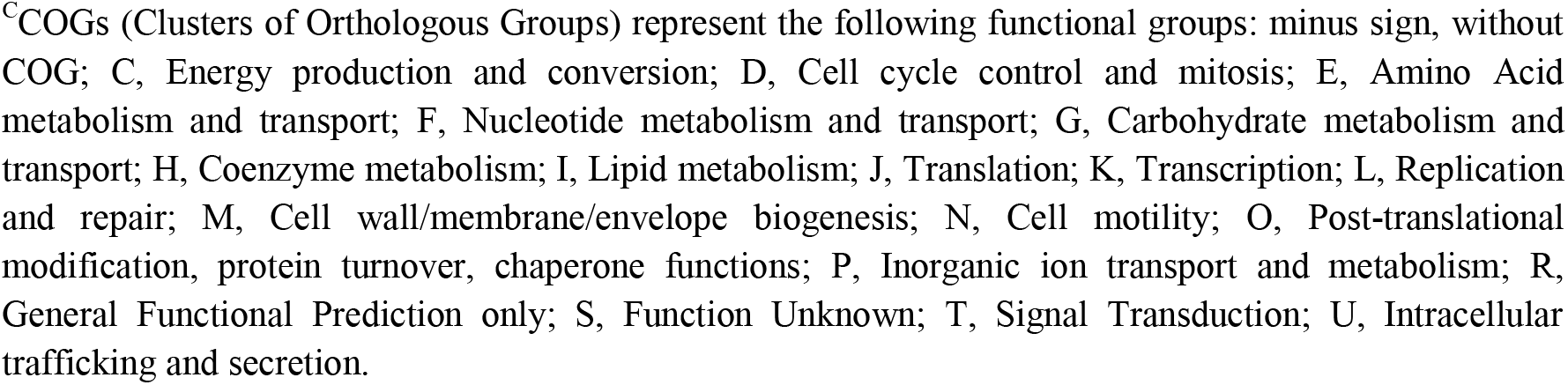
Proteins occurring in *R. etli* OMVs but absent from the periplasm. The presence of the protein at 6 and/or 24 h is indicated by a shaded box.

Previously, we identified 383 extracellular proteins in *R etli* MM culture filtrates (14), which would include OMVs. Ninety of the proteins that occurred exclusively in OMVs or in both periplasm and OMVs (Table 1 and Table S3 (Supplemental material)) were also found in the previously determined exoproteome (14). Thus, nearly one-quarter of the exoproteins identified in our previous study were apparently excreted in OMVs. It should be noted that the mass spectrometric methods used for protein identification in this and our previous (14, 15) work do not allow the quantitation of proteins, but only reveals their presence or absence in a sample.

The 770 proteins identified in the *R. etli* periplasmic and OMV fractions at 6 and/or 24 h (Table S1 in Supplemental Material) represent 12.8 % of the 6022 predicted ORFs encoded in its genome. Only 14.2 % of these proteins are plasmid-encoded, representing less than half of the 32 % of the *R. etli* proteome that is extrachromosomally encoded. There was no significant difference of the relative proportion of plasmid-encoded proteins in the periplasm-only, OMV-only, and in both the periplasm and OMV categories.

Of the 770 proteins identified, 568 and 164 (74 and 21 %) occurred exclusively in the periplasm (Table S1, Supplemental material) and OMV (Table 1) fractions, respectively. Remarkably, only 4.9 % of the total proteins were found in both fractions (Table S3, Supplemental material), which argues against the random inclusion of periplasmic proteins in the OMVs during their formation and supports the idea that specific protein sorting mechanisms are at least partly responsible for determining OMV protein content (30).

The number and identity of periplasmic and OMV proteins produced by bacteria changes with culture age and growth conditions (4,30,33). In *R. etli*, we found significant differences in the identities of the proteins present in the periplasm and OMVs at 6 versus 24 h (Table S1, Supplemental material). In the periplasm-only fraction (Table S2, Supplemental material), 31 and 45 % of the proteins were present only at 6 and 24 h, respectively, and 24 % were present at both 6 and 24 h. For the proteins found exclusively in the OMVs (Table 1), 49 and 42 % were present only at 6 and 24 h, respectively, and 9 % were present at both times. The largely distinct protein profiles for OMVs from log and stationary phase cultures, with relatively few proteins present at both sampling times, indicates a time-dependent differential packaging of proteins into the OMVs. For example, 9 of the OMV-exclusive protein COG groups are comprised of a majority of proteins that are present at 6 but not 24 h. What accounts for the disappearance of these proteins between 6 and 24 h? Possibly, these proteins are selectively degraded within the OMVs, or are released from them, as the culture ages. It has been proposed that different subpopulations of OMVs with a distinctive protein content could exist in the same bacterial culture, but this has hardly been addressed experimentally (35).

Proteins without a dedicated transport mechanism might enter the exoproteome by interacting with one or more proteins that are specifically excreted. We determined potential protein-protein interactions in the *R. etli* proteome using the ProLinks server (http://prl.mbi.ucla.edu/prlbeta/ (36). While highly probable (P = 1.0) interactions were predicted to occur between some members of the total proteome, none were found among the proteins identified in the periplasm and/or OMVs, even at the lowest probability setting (P = 0.4).

### Functional Distribution of Periplasmic and OMV Proteins

An important reason for identifying proteins in the *R. etli* periplasm was to determine if the OMVs also contained a significant number of these proteins. As mentioned, less than 5 % of the proteins identified were shared between the two locations. The identity of periplasmic proteins has perhaps been best established in *E. coli*, which contains a wide functional diversity of proteins among the hundreds that are present (4). The *R. etli* periplasmic proteins are also diverse in functional categories (Fig. 2A). The General functional prediction only COG (R) had the greatest number of proteins (13 % of the total), and proteins involved in amino acid and carbohydrate metabolism and transport were also highly represented. With the exception of the COGs for periplasmic proteins having very few or no members (cell cycle control, replication, motility, secondary metabolism and trafficking/secretion), at least 10 proteins were present in the other COGs and had a relatively even (Fig. 2A) numerical distribution among them.

**Fig. 2.**
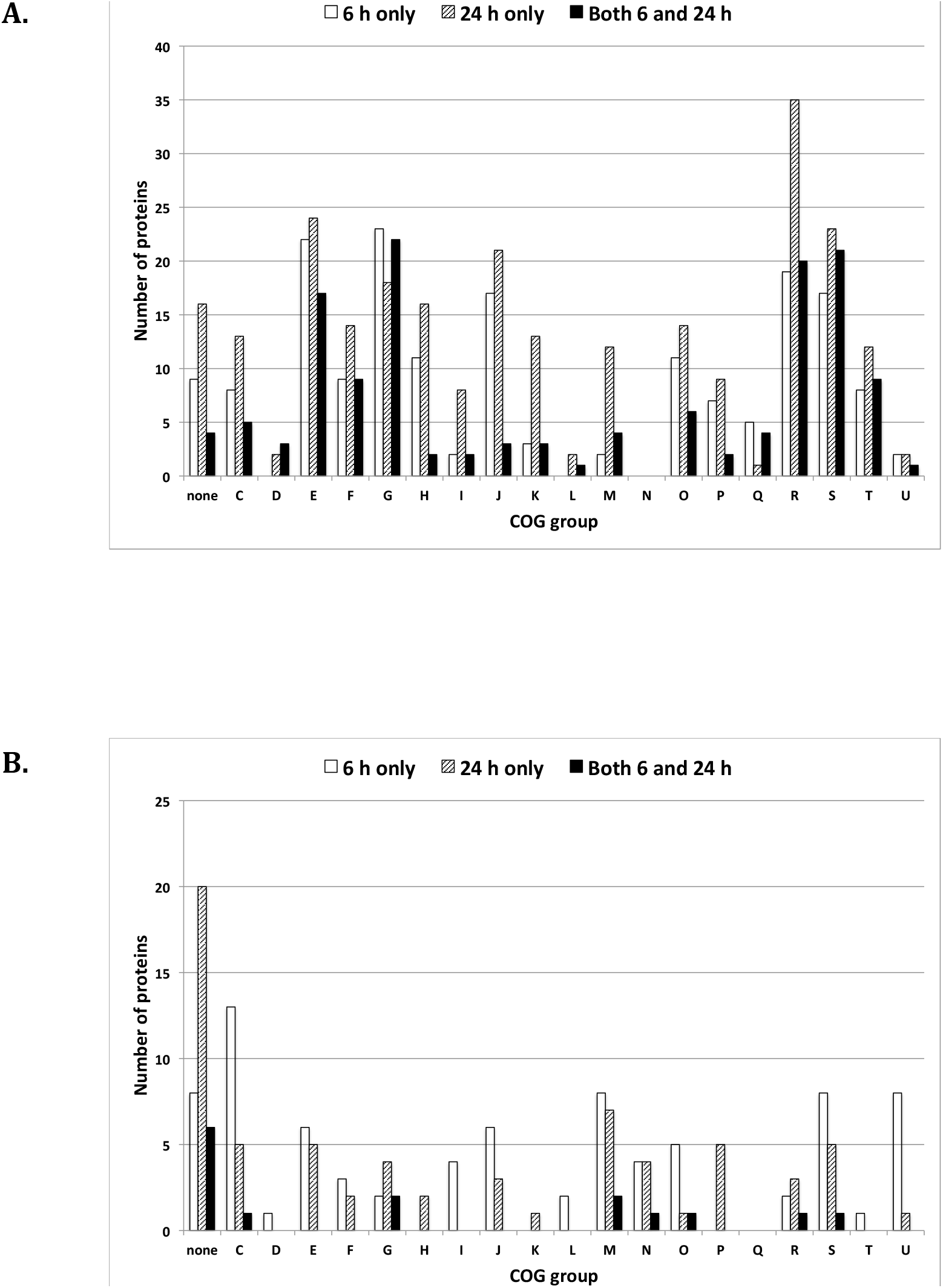
Distribution of periplasm-only (A) and OMV-only proteins (B) by functional category and presence at 6 h, 24 h, or both 6 and 24 h. GOG categories are as described in Table 1.

For proteins present in OMVs but absent in the periplasm, those without an assigned COG are the most abundantly represented category, accounting for about 21 % of the total (Table 1 and Fig. 2B). About 59 % of OMV-exclusive proteins in this COG were present only at 24 h. In comparison, only 5.1 % of present in the periplasm but absent in OMVs are in this category (Fig. 2A). Note that the majority (61-70 %) of the proteins without COG present only in the periplasm or only in OMVs were hypothetical proteins. The rest of the OMV-exclusive proteins were more or less evenly distributed among many of the remaining COGs, with 6 h samples usually having the greatest number of proteins. Over half of the COGs lacked representatives from 6 h and/or 24 h, with several COGs (eg., cell cycle control, coenzyme metabolism, transcription) having a strikingly lower number of total proteins (Fig. 2B). These comparisons highlight the fact that distinct temporal and numerical patterns in periplasm and OMV proteins occur during the growth of *R. etli* in culture. The exoproteins in our dataset (Table S1) were not biased towards being the products of *R. etli* genes expressed as part of specific regulons (37–39) or under conditions of biofilm formation (40).

### OMV-localized proteins of physiological interest

Here we mention some of the OMV proteins with potentially important physiological roles. Based on their analysis of proteins reported in a variety of OMV proteomes, Lee et al (2008) (41) found that many of the proteins belonged to a relatively limited number of protein functional families, including porins, murein hydrolases, multidrug efflux pumps, ABC transporters, proteases/chaperones, adhesins/invasins, and cytoplasmic proteins. Many of the *R. etli* OMV proteins described below fit into one of these categories.

Based on what is known of orthologous proteins in other rhizobia, the Rap proteins and polysaccharidases described below are probably secreted by the *R. etli* PrsDE type I secretion system (T1SS) (42, 43). The 3 autoaggregation/adherence proteins (accessions RHE_RS11065, RHE _RS12180 and RHE _RS23910; Table 1) found in OMVs but not in the periplasm belong to the Rap (Rhizobium adhering proteins) family and include the sole plasmid-encoded Rap paralog and two of the 4 chromosomally encoded Raps found in *R. etli*. These calcium-binding, cell surface-localized proteins are found exclusively in *Rhizobium leguminosarum* biovars and in *R. etli* (44), They are important in rhizobial autoaggregation in both species, and in *R. leguminosarum* bv. trifolii RapA1 influences binding to host cell roots and, when overexpressed, nodulation competitiveness on clover (44–46). Curlin (RHE_RS24865) forms curli fimbriae, surface proteins that are common in bacteria and that in Enterobacteria are important for attachment to host cells and for biofilm formation (47). Whether the *R. etli* Rap proteins and curlin in OMVs acts as an adhesion bridges to surfaces (12) is a topic for future research.

In *R. leguminosarum* bv. viciae, the PlyA, B and C polysaccharidases degrade and reduce the molecular mass of *R. leguminosarum* EPS and also attack carboxymethylcellulose, an analog of plant cell wall cellulose (42, 48). A number of similar polysaccharidases were unique to the *R. etli* OMV fraction, namely the righthanded helix repeat-containing pectin-lyase like PlyB ortholog RHE_RS13345, the contiguously-encoded PlyC ortholog RHE_RS13340 and polysaccharidase RHE_RS03110. In *R. leguminosarum*, polysaccharidases PlyA and PlyB have been well characterized, while the PlyC, which shares a high sequence identity with PlyB has not. PlyA and PlyB are not required for an effective symbiosis between *R. leguminosarum* and pea or vetch (48) but mutants in either polysaccharidase form significantly less biofilm than the parent strain (49). In *R. leguminosarum* PlyA and PlyB are secreted by the TISS and diffuse away from the producing cells, but are inactive until they interact with EPS on the surface of cells in the vicinity. Both enzymes are able to cleave nascent but not mature EPS chains and are not activated by partially-purified *R. leguminosarum* EPS (50). If the *R leguminosarum* polysaccharidases are present in OMVs like their orthologs in *R. etli* (Table 1), it might facilitate their delivery to *R. leguminosarum* cells or host plant roots. That a protein exported by a TISS can be present in OMVs was demonstrated for the *E. coli* α-haemolysin (51).

The *R. etli* OMVs contained the RTX toxin hemolysin-type calcium-binding protein RHE_RS09670. In *R. leguminosarum* bv. viciae, an exported protein of the same family, designated NodO, has been shown to bind calcium and may be important for the attachment of the bacteria to roots (52). Other OMV-localized proteins that determine cell surface characteristics included the glucosyl ketal-pyruvate-transferase RHE_RS16450 (a PssM ortholog), the glycosyl transferase RHE_RS 16485, peptidoglycan-binding protein LysM and the organic solvent tolerance protein OstA. The latter protein has varied roles in membrane synthesis in different bacteria and is an essential protein in *E. coli* (53). The L,D-transpeptidases RHE_RS00275, RHE_RS04095 and RHE_RS06695 likely catalyze alternative peptidoglycan crosslinking reactions. In *E. coli*, the alternative cell wall cross-links introduced by these enzymes are essential for resistance to some antibiotics (54). Antibiotic resistance in rhizobia is potentially important for their competition with antibiotic-producing soil organisms (55). Other cell wall modifying enzymes identified in the OMVs include the peptidoglycan-binding protein RHE_RS09350 and the D-alanyl-D-alanine carboxypeptidase RHE_RS11305.

Many proteins involved in transport or excretion were identified in the OMVs. Three chromosomally-encoded outer membrane *Rhizobium* outer membrane protein A (RopA) orthologs were found in the OMV fraction. The function of these porins in rhizobia is largely undefined but it was determined that the RopA1, RopA2 and RopA3 proteins found in *R. etli* OMVs are not involved in copper transport like the plasmid-encoded RopAe (56), which was not present in the exoproteome. In *Sinorhizobium meliloti* RopA1 is a major phage binding site and, presumably due to other, yet undefined physiological roles, is essential for cell viability (57, 58). Some bacteria produce OMVs containing phage binding proteins as decoy targets for the virus (59, 60). In *R leguminosarum* bv. viciae, RopA1 and RopA2 are secreted, along with polysaccharidases PlyA1 and PlyA2, by the TISS (49). Other secretion-related OMV proteins include the preprotein translocase subunit YajC RHE_RS09360, the ExbD/TolR biopolymer transport family protein RHE_RS17705 and the VirB4 family type IV secretion/conjugal transfer ATPase RHE_RS21955. Two proteins annotated as sugar ABC transporter substrate binding proteins (RHE_RS07970 and RHE_RS16550) have sequence similarities to proteins involved in polysaccharide export. In the 24 h OMVs we identified two IM-localized MexE family multidrug efflux RND (resistance nodulation cell division) transporter periplasmic adaptor subunits, MexE1 and MexE2. These are expected to form part of the HlyD (Type I) multi-drug efflux system. A related protein, RHE_RS18860, encodes an efflux RND transporter periplasmic adaptor subunit. RND efflux pumps contribute to nodulation competitiveness and antimicrobial compound resistance in *S. meliloti* (61). Mex efflux pumps were present in OMVs from *Pseudomonas* species (33). BamA (RHE_RS09805) is the OM component of the ß-barrel assembly machinery (BAM) responsible for the insertion of virtually all OM proteins in Gram-negative bacteria (62). BamD, the outer membrane protein assembly factor that forms part of the Bam complex, was also present in OMVs. Although the reconstituted *E. coli* Bam complex is able to insert proteins into artificial membrane vesicles (63), it is not known whether Bam complex components can do the same in natural OMVs. Porin RHE_RS04090 also has a sequence indicative of an OM beta-barrel protein. Three porins (RHE_RS18285, RHE_RS06895, RHE_RS12455) all predicted to be OM proteins of the porin family common in alphaproteobacteria were present in the OMVs. Nodulation protein NodT RHE_RS17445 is a chromosomally encoded OM lipoprotein that is not involved in Nod factor synthesis or transport. A functional *nodT* is essential for the viability of *R. etli* CE3, where NodT is proposed to play a role in chromosome segregation or maintaining OM stability rather than as an export pump (64). The dicarboxylate transporter RHE_RS15195 (DctA) is expected to be a symbiotically essential gene in *R. etli*, since rhizobial mutants defective in dicarboxylate transport are unable to fix nitrogen (65). Three other ABC transporter substrate binding proteins similar to UgpB are probably involved in glycerol 3-phosphate transport (RHE_RS08805, RHE_RS24975, RHE_RS10590). Polyamines are involved in growth and stress resistance in rhizobia (66), RHE_RS10970 resembles a lysine/arginine/ornithine-binding periplasmic protein that could transport polyamine precursor amino acids into the cell, and RHE_RS22990 is likely a spermidine/putrescine-binding periplasmic protein (PotD). Some bacteria package signal molecules for quorum sensing in OMVs (10, 11, 67). Transporter RHE_RS09265 is a FadL ortholog: in rhizobia, these are important for the uptake of quorum sensing system long-chain *N*-acyl-homoserine lactones. These FadL orthologs are probably involved in transporting long-chain acyl-homoserine lactones across the OM (68), but the presence of a FadL transporter in OMVs could provide for their uptake into vesicles, which could deliver them, perhaps in a concentrated dose, to recipient cells.

Proteins involved in energy production and conversion represent more than 11 % of the OMV-exclusive proteins, over 2-fold more than among the periplasm-only proteins. For oxidative phosphorylation, numerous components of NADH dehydrogenase, cytochrome c, NADH-quinone oxidoreductase and ATP synthase were found principally in the OMVs, although not all of the proteins required to completely assemble these complexes was present.

Among the cytoplasmic proteins present only in OMVs, we found Tme, the NADP^+^-specific malic enzyme that in *S. meliloti* appears to serve as a secondary pathway for pyruvate synthesis during growth on succinate (69). Porphobilinogen synthase is required for the synthesis of tetrapyrrole pigments such as porphyrin and vitamin B12. Ribonuclease PH is involved in tRNA processing. Ribosomal protein subunits accounted for 2.6 and 4.2 %, respectively, of the *R. etli* periplasm - and OM - exclusive proteins and have been found in OMVs from *E. coli* and *Neisseria meningitides* (70, 71).

OMV localized chaperones include the cytoplasmic heat-shock protein Hsp20 and the periplasmic peptidyl-prolyl isomerases SurA and RHE_RS11115, the protease modulators HflC and HflK, and the RNA chaperone Hfq. The latter protein has diverse functions in riboregulation in rhizobia (72).

Finally, because some of the OMV-localized proteins like peptidoglycan modifying enzymes could have antimicrobial activity, we assayed for the ability of purified *R. etli* OMV to inhibit the growth of *Bacillus subtilis* (Fig. S1, Supplemental material). In plate assays, we found that purified OMVs and, especially, ethyl acetate extracts of OMVs, did inhibit the growth of *B. subtilis*. No inhibition was observed with *R. etli* MM culture or culture supernatant, or with ethyl acetate.

In summary, we show here that *R. etli* produces OMVs with significant temporal differences in their protein content during growth in culture. The proteins in the OMVs are largely distinct from those of the periplasmic space and includes many proteins of physiological interest, including some with known symbiotic roles. The excretion of these proteins in OMVs could give a survival or metabolic advantage to free-living or symbiotically-associated *R. etli* cells, and provides an exciting topic of research that we are currently exploring.

## MATERIAL AND METHODS

Bacterial strains and growth conditions. *Rhizobium etli* strain CE3 was maintained in 15 % glycerol stocks prepared from PY rich medium cultures containing 200 and 20 μg/ml of streptomycin and nalidixic acid, respectively. Minimal medium (MM) contained 10 mM each of succinate and ammonium chloride as carbon and nitrogen sources, respectively (14). *Bacillus subtilis* 168 was obtained from E. Martínez (Centro de Ciencias Genómicas-UNAM, Cuernavaca, Mexico) and maintained on PY rich medium (14).

### Isolation of periplasmic proteins

A modification of the hypo-osmotic shock technique used with *R leguminosarum* by Krehenbrink *et al* (19) was used to isolate *R etli* periplasmic proteins. To do this, bacterial cells obtained by centrifugation (6000 x g) were washed twice with cold 1.0 M NaCl, resuspended in cold sucrose buffer (20% (w/v) sucrose, 1 mM EDTA and 10 μl of protease inhibitor cocktail (#13743200, Roche) per 10 ml and incubated for 5 minutes at room temperature. The samples were centrifuged at 6000 x *g* for 10 minutes at 4°C, the pellets were resuspended in cold distilled water for 5 minutes at room temperature followed by centrifugation as above for 30 minutes. The supernatants were transferred to new tubes and precipitated with 2.5 volumes of cold acetone followed by overnight incubation at -20°C. The samples were centrifuged at 8000 x *g* for 45 minutes, the pellets were washed with 80% acetone and resuspended in urea solubilization buffer (7 M urea, 2% (v/v) CHAPS, 1 mM DTT) and stored at -20°C until analysis.

### Isolation of OMV proteins

Cultures of *R. etli* were grown for 6 and 24 h in 2 L of MM under the conditions described previously (14). To isolate OMVs, the cultures were centrifuged at 6000 x *g* for 10 minutes at 4°C. The supernatant was filtered through a 0.22 μm filter and concentrated to dryness by lyophilization. Lyophilized samples were resuspended in 1 mM Tris-HCl (pH 8.0) buffer and ultracentrifuged in transparent tubes (Beckman 14 X 89 mm), at 100,000 x *g* for 2 h at 4°C in a SW41 swinging bucket rotor (Beckman). The pellet was resuspended in 500 μl of 1 mM Tris-HCl (pH 8.0 buffer), the proteins were extracted with phenol and kept at -20°C until use (15).

### Transmission electron microscopy

Samples from different purification stages of the of OMV and periplasm isolations were resuspended in 20 mM Tris-HCl (pH 8). Five μl of purified OMV samples were applied to 400-mesh copper grids, 2% acid phosphotungstic was added followed by incubation for 1 min at room temperature. Grids were observed in a JEM1011 (JEOL, Japan) at 100 kilowatts of acceleration voltage.

### SDS-PAGE and protein digestion

The protein concentration of OMVs and periplasm samples was determined by the Bradford method (73). Gel electrophoresis was carried out on 12.5% resolving gels loaded with 20 μg of total protein per lane. Gels were stained with Coomassie blue and 5 mm wide gel slices were excised, transferred to microcentrifuge tubes and covered with 20 μl of 20% EtOH. Gel slices were sequentially washed with 100mM Tris-HCl (pH 8) and 50 % aqueous acetonitrile, reduced with 50 mM DTT in 50 mM Tris-HCl (pH 8) for 30 min at 37°C. Samples were alkylated with 50 mM iodoacetamide in 50mM Tris-HCl (pH 8) for 30 min at 37°C (in darkness), and incubated for 5 h at 37°C with trypsin (10 ng/μl, sequencing grade, Promega, Switzerland) in 20 mM Tris-HCl (pH 8.). The tryptic fragments were extracted with 20 μl of 20% (v/v) formic acid. Two independent experiments to obtain OMV and periplasmic fractions were performed, with 85% of the exoproteins identified being found in both experiments. Only these proteins are included in the data reported here.

### Mass spectrometry

Peptide sequencing was performed on a LTQ XL-Orbitrap mass spectrometer (Thermo Fisher Scientific, Bremen; Germany) equipped with a Rheos Allegro nano flow system with AFM flow splitting (Flux Instruments, Reinach; Switzerland) and a nano electrospray ion source operated at a voltage of 1.6kV. Peptide separation was performed on a magic C18 nano column (5 μm, 100 Å, 0.075 x 70 mm) using a flow rate of 400 nl/min and a linear gradient (60 min) from 5 to 40 % acetonitrile in H_2_O containing 0.1% formic acid. Data acquisition was in data dependent mode on top five peaks with an exclusion for 15 sec. Survey full scan MS spectra were from 300 to 1800m/z, with resolution R=60000 at 400 m/z, and fragmentation was achieved by collision induced dissociation with helium gas in the LTQ.

### Protein identification and prediction of subcellular localization

Mascot generic files (mgf) were created by means of a pearl script using Hardklor software, v1.25 (M. Hoopmann and M. MacCoss, University of Washington). MS/MS data (mgf files) were submitted to EasyProt (version 2.3) for a search against the SwissProt database (Rhizobium_Homo_Tryp_ForRev (20100415) in two rounds. First round parameters were: parent error tolerance 20 ppm, normal cleavage mode with 1 missed cleavage, allowed amino acid modifications (fixed Cys_CAM, variable Oxidation_M), minimal peptide z-score 5, max p-value 0.01 and AC score of 5. Second round parameters were: parent error tolerance 20ppm, half cleaved mode with 4 missed cleavages, allowed amino acid modifications (variable Cys_CAM, variable Deamid, variable phos, variable Oxidation_M, variable pyrr) minimal peptide z-score 5, max p-value 0.01. Protein identifications were only accepted with an AC score of 10, meaning when two different peptide sequences could be matched. The PSLpred program (25) was used to predict the subcellular localization of the proteins. We determined potential protein-protein interactions among the *R. etli* exoproteome using the ProLinks server (http://prl.mbi.ucla.edu/prlbeta/ (36).

### Cytotoxic Activity of OMVs

OMVs purified from 24 h *R. etli* CE3 were extracted with ethyl acetate. *B. subtilis* 168 was cultured overnight in PY at 30°C, cells were washed twice with sterile distilled water and diluted with sterile water to an optical density of 0.05 at 540 nm. Two-hundred μl of the *B. subtilis* cells were plated on petri dishes of PY supplemented with 0.1 mM CaCl_2_. Whatman filter paper circles 0.5 cm in diameter were placed on the plate and impregnated with 5 μl test samples. The plates were incubated for 48 hours at 30°C before determining the presence of zones of growth inhibition surrounding the paper discs.

## SUPPLEMENTAL MATERIAL

Supplemental material for this article may be found at https://doi.org/……AEM….

## ACKNOWLEDGMENTS

We thank Esperanza Martínez (CCG-UNAM) for supplying *B. subtilis* 168. Part of this work was supported by CONACyT grant 220790 and DGAPA-PAPIIT grant IN213216.

---

Fig. S1. Growth inhibition of *Bacillus subtilis* 168 by OMVs preparations from *Rhizobium etli* CE3. The filter paper discs on the lawn of *B. subtilis* cells were treated with 1) Strain CE3 culture; 2) Unfiltered culture supernatant; 3) Filtered (0.22 μM) culture supernatant; 4) Ethyl acetate extract of OMVs; 5) Purified OMVs; 6) Ethyl acetate.

